# Bi-clustering based biological and clinical characterization of colorectal cancer in complementary to CMS classification

**DOI:** 10.1101/508275

**Authors:** Sha Cao, Wennan Chang, Changlin Wan, Yong Zang, Jing Zhao, Jian Chen, Bo Li, Qin Ma, Chi Zhang

**Author notes:** To whom correspondence should be addressed. Correspondence is also addressed to Sha Cao and Qin Ma.

## Abstract

In light of the marked differences in the intrinsic biological underpinnings and prognostic outcomes among different subtypes, Consensus Molecular Subtype (CMS) classification provides a new taxonomy of colorectal cancer (CRC) solely based on transcriptomics data and has been accepted as a standard rule for CRC stratification. Even though CMS was built on highly cancer relevant features, it suffers from limitations in capturing the promiscuous mechanisms in a clinical setting. There are at least two facts about using transcriptomic data for prognosis prediction: the engagement of genes or pathways that execute the clinical response pathway are highly dynamic and interactive with others; and a predefined patient stratification not only largely decrease the statistical analysis power, but also excludes the fact that clusters of patients that confer similar clinical outcomes may or may not overlap with a pre-defined subgrouping. To enable a flexible and prospective stratified exploration, we here present a novel computational framework based on bi-clustering aiming to identify gene regulatory mechanisms associated with various biological, clinical and drug-resistance features, with full recognition of the transiency of transcriptional regulation and complicacies of patients’ subgrouping with regards to different biological and clinical settings. Our analysis on multiple large scale CRC transcriptomics data sets using a bi-clustering based formulation suggests that the detected local low rank modules can not only generate new biological understanding coherent to CMS stratification, but also identify predictive markers for prognosis that are general to CRC or CMS dependent, as well as novel alternative drug resistance mechanisms. Our key results include: (1) a comprehensive annotation of the local low rank module landscape of CRC; (2) a mechanistic relationship between different clinical subtypes and outcomes, as well as their characteristic biological underpinnings, visible through a novel consensus map; and (3) a few (novel) resistance mechanisms of Oxaliplatin, 5-Fluorouracil, and the FOLFOX therapy are revealed, some of which are validated on independent datasets.

## INTRODUCTION

Colorectal cancer is the fourth most frequent cancer in the United States, which accounts for more than 8% of adult cancer incidence and 8% cancer deaths in 2018 (1). Epidemiology data suggests the average five-year survival rate of CRC is 64.9%, while more than 80% of patients die from the disease in five years in the case of metastasis (2, 3). Amongst all, intra-tumor heterogeneity could account for a significant part of poor treatment response. CRC is one of the cancer types with most clearly delineated heterogeneity, a few molecular subtyping methods have been developed, with the goal that it will facilitate the translation of molecular subtypes into the clinic (4–12). Among these, the Consensus Molecular Subtype (CMS) classification has been accepted as a standard practice for colorectal cancer (CRC) stratification (4, 5). CMS classification was derived from a cohort of 18 independent gene expression data sets with 4,151 samples of CRC, and it has stratified more than 85% of these CRC samples into four classes with distinct molecular features and prognoses (4). However, to the best of our knowledge, it remains largely undiscovered regarding the CMS class specific prognosis and predictive gene markers and relevant biological underpinnings, and further class based targeted interventions (4). A major challenge for identification of disease subtype specific biomarkers is that the statistical power will be largely reduced once the analysis is restricted to a pre-defined stratification. This preprocessing is only meaningful when the stratification perfectly aligns with the diversity among samples in response to the prospective clinical outcome. Otherwise, the pre-stratification would severely limit our power in identifying novel alternative mechanisms underlying the clinical outcomes. These largely undermine the practicality of the CMS classification, and limited its capacity for clinical translation.

It is imperative to develop a framework that enables us to study the possible alternative regulatory mechanisms in cancer in recognition of the patients’ heterogeneity. We utilized a non-parametric approach to identify gene expression modules pertinent to sub-populations, namely, bi-clustering. Bi-clustering analysis is a technique to identify gene co-expression structures specific to certain and sometimes to-be-identified subsets of samples (13, 14). The algorithm outputs data blocks, each containing subset of samples and features in a sub-matrix format, called bi-clusters (BC). We have recently released a new bi-clustering R package QUBIC-R, which enables identification of bi-clusters (BCs) in whole-genome level transcriptomics data set and has been shown to have competitive performance compared with others (15–17). We investigated the identified BCs from a large collection of gene expression data of CRC to: (1) identify potential gene modules specific to a subset of CRC samples; (2) provide a mechanistic interpretation of the CRC subtypes, in retrospective of CMS in particular; and (3) identify prognosis markers and alterative drug resistance mechanisms specific to different disease subtypes. Under the bi-clustering framework, where there is no need of pre-defined stratification, we have the power to analyze the data as an intact entity. Each BC potentially contains signature and coherent gene modules existent in a subgroup of patients, that reflects the heterogeneous gene expression patterns between samples within and out of the BC. The gene subsets may enrich certain biological pathways that could lead to substantially deeper biological understanding for molecular stratification of CRC. More importantly, any existing sub-grouping methods, such as CMS, could be studied and integrated with the produced BCs retrospectively.

Thus, we believe our computational framework based on bi-clustering provides a powerful tool for systematic interrogation of the disease in different clinical settings without compromising the analysis power. The analysis fully recognizes the large heterogeneity within CRC patients, some of which may be strongly associated with existing CRC sub-classes defined by various clinical and genomic features, while the rest will provide novel alternative ways for us to better understand the disease. Our key results include: (1) a comprehensive annotation of the local low rank module landscape of CRC; (2) a novel consensus map demonstrates that CMS IV seem to resemble a mixture of CMS I-III with high stromal infiltration, while CMS I-III also show characteristics of other classes; (3) disease progression free survival of CRC are largely determined by micro-environmental alterations while the overall survival is more associated with the level of stromal infiltration in a CMS dependent manner; and (4) a few (novel) resistance mechanisms of Oxaliplatin, 5-Fluorouracil, and the FOLFOX therapy are revealed, some of which are validated on independent datasets.

## RESULTS

In this study, we conducted a bi-clustering analysis in multiple large CRC data sets aiming to: (1) generate a comprehensive annotation for the landscape of coherent co-expression modules specific to different subsets of samples; (2) identify CMS class dependent BCs and annotate biological mechanisms of the BCs and CMS class, (3) identify prognosis predictive BC that are CMS class dependent/independent; (4) identify alternative drug resistance mechanisms. By applying our in-house algorithm QUBIC-R on eight colon cancer transcriptomics data sets with 1,440 samples, we have identified ~4,000 significant BCs on average in each data set (Table 1). Each of the BC is further annotated by its statistical significance, the pathways enriched by its genes, and the associations of its samples with CMS class, clinical features, and patients’ survival.

**Table 1.** Bi-clustering information of the eight data sets

| Data ID | #Identified BCs | #Significant BCs | #Pathway enriched BCs | #CMS BCs | #DFS and OS BCs | #Other clinically associated BCs |
| --- | --- | --- | --- | --- | --- | --- |
| **GSE14333** | 9631 | 6547 | 2597(39.7%) | 2512(38.4%) | 448(6.8%) | 452(6.9%) |
| **GSE17536** | 11255 | 4806 | 2187(45.5%) | 1425(29.7%) | 284(5.9%) | 63(1.3%) |
| **GSE29621** | 8167 | 1758 | 582(33.1%) | 289(16.4%) | 73(4.2%) | 56(3.2%) |
| **GSE33113** | 9238 | 2836 | 795(28%) | 958(33.8%) | 136(4.8%) | 3(0.1%) |
| **GSE37892** | 10644 | 4452 | 1600(35.9%) | 1202(27%) | 130(2.9%) | 101(2.3%) |
| **GSE38832** | 5845 | 4319 | 2603(60.3%) | 1705(39.5%) | 335(7.8%) | 0(0%) |
| **GSE39582** | 8267 | 4658 | 1200(25.8%) | 2894(62.1%) | 1068(22.9%) | 1847(39.7%) |
| **TCGA_COAD** | 2697 | 2632 | 1077(40.9%) | 743(28.2%) | 183(7%) | 954(36.2%) |

### Analysis Pipeline and Statistical Consideration

Figure 1A shows the analysis pipeline of this study. Gene expression profile of each data set is first discretized to a binary matrix in preparation for the bi-clustering analysis. Figure 1B details the bi-clustering analysis procedure. For each gene and an integer K, expression profile of the gene was non-parametrically discretized to generate K binary vectors, where 1s represent those samples having the gene’s expression in the 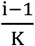 to 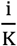 quantile in the ith vector, i=1,…,K. Otherwise, the vectors have zero values. In this way, the original m × n gene expression matrix with m genes and n samples is expanded to a Km × n binary matrix, as shown in Figure 1B and detailed in Methods section. Then, submatrices enriched by 1s in the discretized matrix are identified as BCs heuristically. Obviously, small K would blur the variability of gene expression across samples, and large K would severely undercut the power of bi-clustering and result in small “narrow” bi-clusters. We also noticed that the proportion of the largest subtype in CRC is about 1/3, and after testing K=2, 3, 4, and 5, we found that the discretization with K=3 results in largest number of significant associations between BCs and biological and clinical features (see details in Methods and Supplementary Figure S1). Considering these, K=3 is selected for all future analysis. Each identified BC consists of a subset of samples and a group of genes, in which the genes are consistently expressed highly, moderately, or lowly over the subset of the samples, forming a tight rank-1 co-expression module specific to these samples. We utilized a rigorous assessment method for the statistical significance test of the BC’s (details in Methods section), and those significant BCs are further examined to see whether genes in a BC enrich a certain pathway or gene set, and samples in a BC significantly over-represent a certain phenotype. The analysis pipeline is implemented with our newest QUBIC-R package, which was recently optimized for large-scale matrices (15).

**Figure 1.**
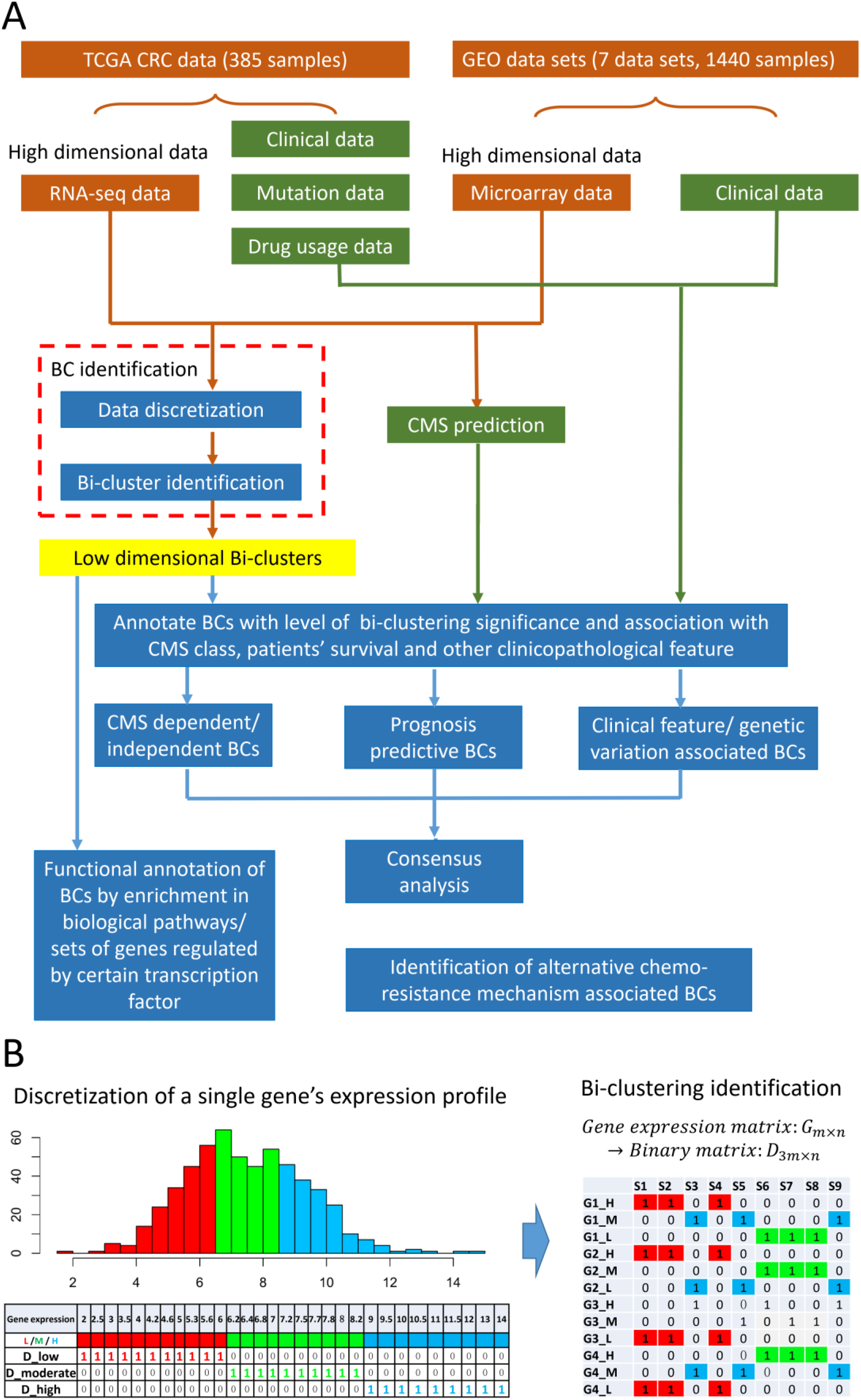
**(A) General analysis pipeline.** The analysis was conducted to one TCGA RNA-seq and seven microarray datasets. BC identification of each high-dimensional data sets is conducted by a discretization followed by a bi-cluster identification step (see details in B). The identified BCs are further annotated by their associations with biological pathways, CMS class, and patients clinical and prognostic features. Consensus analysis of the BCs throughout multiple data sets was further conducted. BCs were further associated with response to different chemo-drugs for identification of alternative chemo-resistance mechanisms. **(B) Data discretization and bi-clustering procedures.** The histogram on the left illustrates the distribution of a gene’s expression. The gene expression is represented as three 0-1 vectors (D_high, D_moderate and D_low), corresponding to samples with top (blue), medium (green) and bottom (red) 1/3 expression level of the gene, respectively. The discretized data are then merged together that expand an original m × n gene expression matrix to a 3m × n binary matrix, as shown in the right panel. BCs enriched by 1s are further identified by QUBIC-R.

Features/outcomes that are of particular interests in this study include: 29 clinical features/outcomes in supplementary Table 1; 73 cancer-associated gene mutations (supplementary Table 1); and treatment responses to three chemo therapeutic drugs namely 5-Fluorouracil, Oxaliplatin, and the combination of 5-Fluorouracil, Oxaliplatin and Leucovorin. Functional annotation of the BCs are conducted against 1329 pathways and gene sets in Msigdb (18). The analysis was applied to transcriptomic data of 1,440 patient-derived CRC tissue samples including the TCGA COAD RNA-Seq data set, as well as seven microarray data sets (GSE14333, GSE17536, GSE29621, GSE33113, GSE37892, GSE383832 and GSE39582) measured by Affymetrix UA133 plus 2.0 array platform. (See detailed data information in Method). The computational pipeline and key statistics of this work is provided in GitHub via https://github.com/changwn/BC-CRC, which can be readily transplanted for similar analyzes in other disease scenarios. All the supplementary files could be found in the same GitHub space.

### Comprehensive association studies of BCs with functional gene sets and various clinical/biological features

A total of 65,744 BCs are identified in the eight primarily analyzed data sets, and on average, ~4,000 BCs are found to be significant in each data set (Table 1). Complete gene/sample information of all the significant BCs are provided for each dataset via R data space through the GitHub link, with a description listed in Supplementary table 2. For each significant BC, we comprehensively investigated whether: (1) genes in the BC significantly enrich biological pathways or gene sets (p<1e-6); (2) samples in the BC are significantly associated with CMS class (p<0.005); (3) samples in the BC are significantly associated with clinical features such as age, gender, races and pathological stages (p<0.005); (4) samples in the BC are significantly associated with prognostic outcomes, namely patients’ overall and disease free survival (p<0.005); (5) samples in the BC are significantly associated with genomic mutation profiles (p<0.005); and (6) samples in the BC are significantly associated with the response to three selected chemo-drugs (p<0.005). Figure 2A shows the proportion of BCs with significant annotations of the first four types of associations in the eight data sets. On average, 71.79% (22,981/32,008) of the significant BCs can be significantly annotated by at least one of the four associations in the eight data sets, with detailed numbers listed in Table 1. Complete annotation of the BCs is also provided through GitHub and described in Supplementary Table 2. Note that (5) and (6) are specific to TCGA-COAD dataset. We will discuss (6) in more details in a separate section. Results for additional clinical features, such as TNM stages, not present in all datasets, together with (5), are all listed in supplementary Table 1.

**Figure 2.**
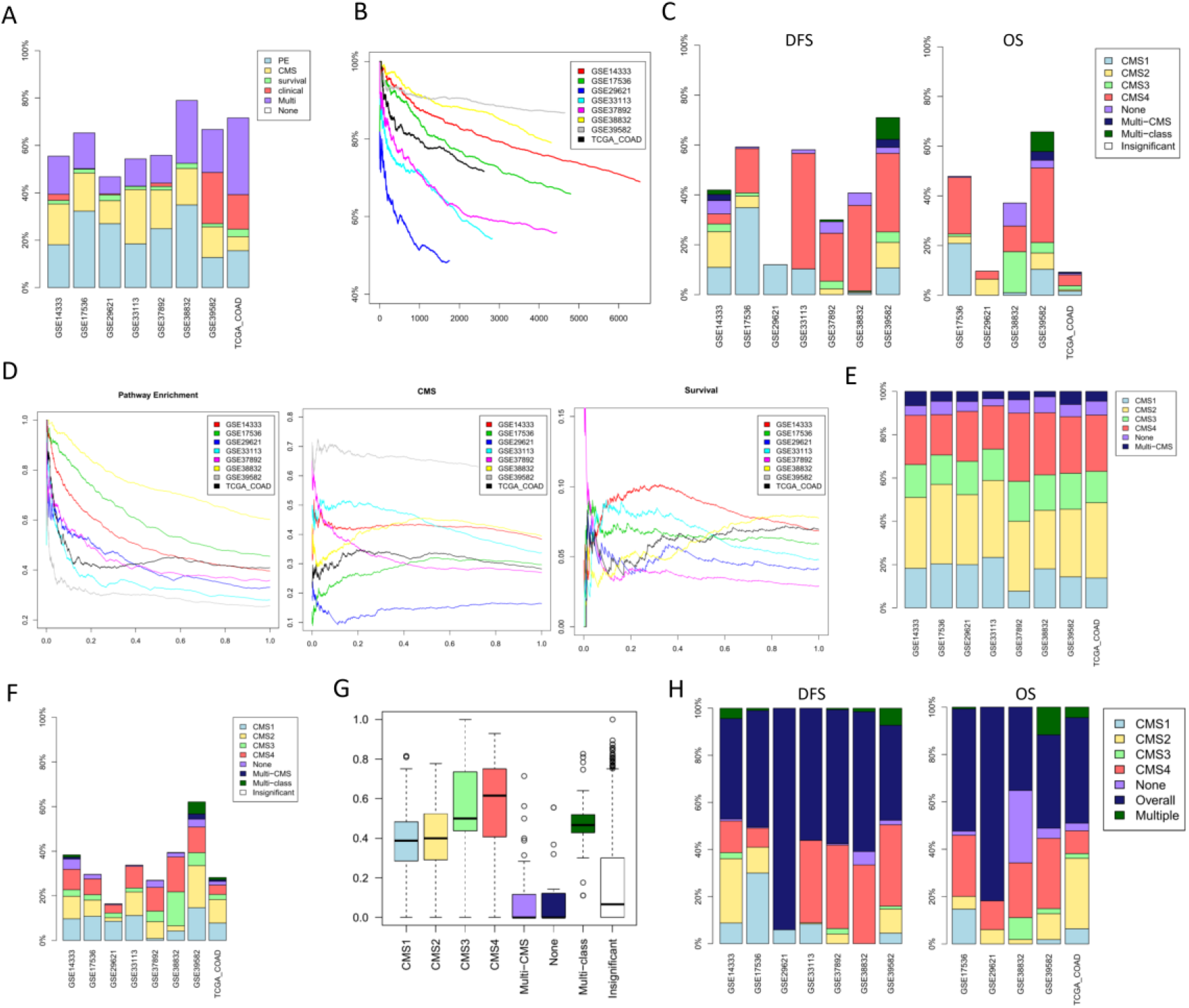
Statistics of the BC landscape in the eight data sets. (A) Proportions of the BCs (y-axis) associated with biological pathways (PE), CMS, patients’ DFS/OS survival, clinical features, and their combinations (Multi) in each data set (x-axis). (B) Cumulative rates of BCs (y-axis) with at least one of the four types of annotations versus ranks of BCs (x-axis). The BCs are ordered by their bi-clustering significance in a descending manner in each data set. (C) Proportions of the BCs (y-axis) that are associated with certain CMS classes among the BCs with significant associations to patients’ survival, including DFS and OS, in each dataset (x-axis). (D) Cumulative rates of BCs (y-axis) significantly associated with biological pathways (left), CMS classes (middle) and patient’s DFS and OS (right) versus the quantiles of the bi-clustering significance (x-axis). For example, a “0.2” quantile means the top 20% significant BCs. (E) Proportions of the BCs (y-axis) with significant associations to different CMS classes in each data set (x-axis). (F) Among the BCs with significant associations to patients’ survival, the proportions of the BCs (y-axis) associated with CMS types in each data set (x-axis). (G) For BCs associated with different CMS class, the average overlapping rates (y-axis) between the genes in the BC and CMS marker genes in each dataset (x-axis). (H) Among all the DFS/OS associated BCs, the proportion of the BCs (y-axis) that significantly over-represent a (sub)sample class in each dataset (x-axis). In (C), (E) and (F): None: CMS unclassified samples; Multi-CMS: a class of samples falling into more than one CMS classes; Multi-class: a class of BCs significantly associated with more than one CMS classes. In (H): None: CMS unclassified samples; overall: the BCs associated with survival throughout all patients, but not with a particular CMS class; Multiple: the BCs associated with patients’ survival specific to the patients of more than CMS classes.

Figure 2B shows the cumulative ratio of the BCs that show significant annotations for at least once, among pathway, CMS class, patients’ prognosis and other clinical outcomes (y-axis), wherein the BCs are ordered by their bi-clustering significance levels on a descending order (x-axis). On average, more than 80.7% of the top 20% significant BCs and 66.4% of all significant BCs could be significantly annotated in the eight data sets, indicating more significant BCs tend to be more biologically/clinically relevant. This shows that our bi-clustering algorithm could indeed identify local modules that bear biological/clinical significance. In general, for the most significant BCs (p<1e-200), their genes tend to have strong associations to biological pathways, including cell cycle, cell proliferation, cell death, biosynthesis and metabolism of nucleic acid, mRNA and protein, cytoskeleton synthesis, protein phosphorylation, cell membrane, cell adhesion, and immune response and chemokine activity pathways (Figure 3). However, their samples don’t seem to be significantly associated with existing clinical features or CMS classes, meaning that these BCs may be general to the large population. In the next level (1e-200<p<1e-50), the BCs associated with CMS class or other clinical features are with relatively smaller sizes and less significance compared to the first level, and these BCs enrich a different group of biological pathways including immune response, extracellular matrix, cytoplasmic part, O linked and N linked protein amino acid glycosylation, cell membrane, protein modification, lipoprotein biosynthesis and lipid metabolism, ABC transporter, steroid hormone metabolism and signaling pathway.

**Figure 3.**
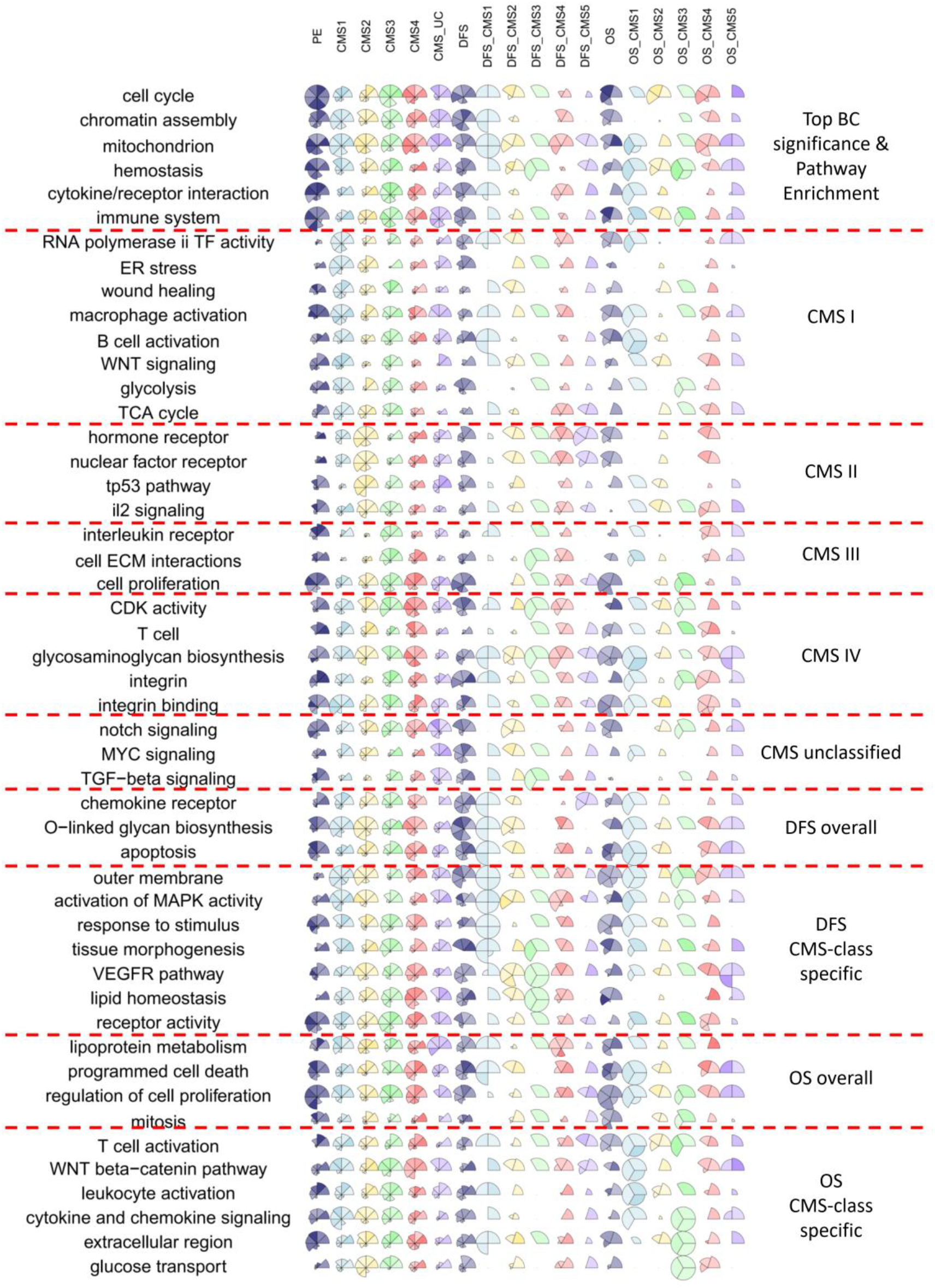
A consensus map representing the intricate relationships between key pathways and clinical features, as well as similarities among different clinical features. The top of the figure shows 18 different pools that the BCs in each dataset are placed in, and the left of the figure shows the pathways that are consistently enriched by the BCs in the pool. Each pie graph consists of up to eight sectors, one sector for one dataset, depending on whether DFS or OS data is available for the dataset. For each of the 18 pools in each dataset, only the BC with the highest enrichment significance of the pathway is selected, and the level of enrichment is presented by the radius and shade of the sectors: the larger the radius, the more the genes in the corresponding pathway are being hit the BC; the darker the shade, the more significant the enrichment is using genes in the BC for the pathway. On the right, the pathways are re-grouped into 10 categories, so that the pathway is assigned to the group that it most significantly represents.

On average, we have seen that 44.7% of the DFS associated and 33.9% of the OS associated BCs are also associated with at least one CMS class while the rest are CMS classification independent, as shown in Figure 2C, suggesting possible CMS class specific prognosis markers. Most of the CMS dependent DFS associated BCs are associated with CMS class I and IV while some OS associated BCs were found to be independent of the CMS classes.

The ratio of BCs that are significantly associated with biological pathways (left), CMS classes (middle) and patient’s DFS and OS (right) versus the quantiles of the bi-clustering significance are shown in Figure 2D. Again, we observe that the more significant BCs tend to significantly enrich more biological pathways. Similar patterns are not identified for CMS class in all datasets. This coincides with our initial motivation that: patients stratifications should not be fixed for all the clinical/biological outcomes, as each of them may have different levels of diversity, and even the most cancer relevant stratification, such as CMS, may not perfectly align with the true subtypes with regard to a certain prospective outcome. Interestingly, BCs associated with patients’ survival, including DFS and OS, fall into two groups: one group accounts for ~30% of the DFS/OS associated BCs with higher significance (p~1e-200<p<1e-80), which shows an overall significant association with DFS/OS regardless of CMS. BCs in this group enrich a diverse set of signaling transduction pathways including NOTCH, RHO factor, TRKA receptor, EGF, RAS, cell surface/ kinase receptor, glycoprotein, chemokine and other immune response related signaling pathways. The other group is formed by BCs with relatively lower significance (p~1e-80<p<1e-20), and their associations with DFS/OS tend to exhibit CMS dependency. This means that the DFS/OS associations are diverse among CMS classifications. Biological characteristics of these BCs are discussed in the following sections.

### A consensus functional annotation of the bi-cluster landscape

Our analysis has revealed that BCs associated with different clinical features enrich distinct sets of pathways, suggesting that different biological/clinical features are characterized by different responsive mechanisms. Among these BCs, a notable portion exhibit a CMS-dependent manner. To help us better understand the functional annotations of these BCs, and the underlying sub-groupings they may represent, we summarized the biological pathways that are consistently enriched by the BCs across all datasets, that do show significant signs of clinical associations, including CMS, OS, DFS and their intersections. We call this a consensus functional annotation of the BC landscape in CRC. As shown in Figure 3, 49 pathways/gene sets in total are examined, and here is how these pathways were selected. We first placed the BCs of each dataset into 18 pools shown on the top of the figure: BCs of top bi-clustering significance, over-representing CMS I, II, III, IV, unclassified, associated with DFS in general, associated with DFS and CMS I, II, III, IV, unclassified, associated with OS in general, associated with OS and CMS I, II, III, IV, unclassified, for each dataset. For each BC in each pool of each dataset, pathway/gene set enrichment was performed, and within each pool, the pathways that are enriched most consistently across all datasets are selected, as shown on the left of the figure. This results in a subset of pathways/gene sets that are consistently enriched by BCs that are shown to have one of the 18 characteristics.

To have an even finer view, we drew pie graphs with sectors of varied radius and shade to provide a more quantitative measure of the intricate relationships among pathways/gene sets and different phenotypes. Each pie graph consists of up to eight sectors, one sector for one dataset, depending on whether DFS or OS data is available for the dataset. The radius of the sector shows the proportion of genes in the pathway that are hit by the BC in the dataset, and shade of the sector shows the significance of the enrichment test for the genes in the BC against the pathway. The larger the radius, the more the genes are being hit the BC; the darker the shade, the more significant the enrichment is. Note that for each pool, only the BC with the highest enrichment significance of the pathway is selected, in drawing the radius and shade of the sectors. Details of the color parameters are shown in Supplementary Note.

Moreover, to exhibit how similar the 18 different pools or phenotypes are, we regrouped the 47 pathways, and found that they in fact fall into 10 categories: BCs of top bi-clustering significance, over-representing CMS I, II, III, IV, unclassified, associated with DFS, associated with DFS in a CMS dependent manner, associated with OS, associated with OS in a CMS dependent manner, as shown on the right of the figure. The re-grouping was done in such a way that each pathway was given a score based on the average radius and shade of the pie graph over all datasets, namely, the hitting frequency and the enrichment significance value, and was then assigned to one of the 10 categories with a highest score. The 10 categories we used here are very similar to the 18 characteristics or pools we presented earlier, only in a coarser way.

This consensus map is a novel visualization that greatly helps us visualize for samples in different cancer subtypes and key clinical outcomes, how they express distinct functional pathways, and they relate to each other and to what extent they resemble, and the resolution is for each pathway and each dataset. As shown in Figure 3, for samples in different CMS classes, they are characterized by different pathways/gene sets: CMS I by ER stress, wound healing, macrophage and B cell activation, WNT signaling and glucose metabolisms; CMS II by hormone receptor, TP53 and IL2 signaling; CMS III by cell proliferation and cell matrix-adhesion; CMS IV by T cell activation and Cyclin dependent kinase; and unclassified samples by notch, MYC and TGF-beta signaling pathways. Moreover, different CMS classes don’t seem to be completely isolated. BCs associated with CMS I are also enriched by immune signaling pathways including IL-3, -5, -6, -12, -27, STAT, and interferon gamma signaling pathways, as well as nucleotide biosynthesis, WNT signaling, lipid metabolism, and glycolysis pathways, which are markers of CMS II and III classes (4). Considering that CMS I is a subtype with high MSI and strong immune cell activation (4), our observation clearly suggests that there are distinct subgroups inside CMS I class with different immune activation status that display CMS II-like characteristics with high expression of epithelial and WNT signaling markers and CMS III-like characteristics of metabolism dysregulations. More intriguingly, the BCs associated with CMS class IV fall into two categories: one enriched by integrin binding, epithelial cell cycle, cell death, cell-cell and cell-matrix adhesions pathways, while the other enriched by immune response, MYC and WNT signaling, and metabolism pathways. The first category show expression of cancer and stromal cell marker genes, suggesting different levels of stromal cell infiltration in CMS IV class samples. In contrast, the second category enriches marker genes of CMS class I-III, suggesting there are subgroups of CMS IV samples with distinct characteristics of CMS class I, II or III. CMS IV is a subtype with high stromal infiltration and angiogenesis (4). Our previous study has identified a dynamic population of mesenchymal-like cells with similar markers as CMS IV (19). With these observations, we suspect that CMS IV is a combination of CMS I-III but with higher proportion of stromal cells, hence higher expression of mesenchymal cell markers and lower rate of somatic mutations. However, it is noteworthy that the CMS IV cancers have generally poorer prognosis comparing to CMS I-III, indicating the level of stromal infiltration may serve as an important prognosis marker for all the CMS classes. We have also seen that a number of BCs associated with CMS II and CMS III are enriched by marker genes of other CMS classes. The BCs associated with the unclassified samples are enriched by signaling pathways of MAPK, P38, GPCR, NOTCH, TGF-beta, ARF6 and other kinase receptors and pathways responsive to micro-environment stresses including ER stress, oxidative stress, dysregulated immune activation and extracellular matrix malfunction. We suspect that these samples are with activation of specific signaling pathways or with distinct micro-environment stresses that cause varied gene expressions, hence cannot be classified by the distance based CMS classifier. A consensus functional annotation of the BCs enriching different CMS classes are given in Supplementary Table 3.

### Heterogeneous prognosis of CRC in retrospective of CMS

For all the eight data sets, on average 19.2% (12,641/65,744) of the BCs are significantly associated (p<0.005) with at least one of the CMS classes, and among these, the proportion of BCs associated with each class is shown in Figure 2E. On average, BCs associated with at least one CMS class only cover 23.6%, 15.6%, 30.1% and 24.1% of the samples for CMS I-IV, respectively (shown in Supplementary Figure 2), suggesting that most of the underlying cancer sub-groups may not align perfectly well with the CMS classification. Comparing the proportion of samples in the BCs falling under different CMS class (shown in Figure 2F), there are relatively more BCs aligning with CMS class I and IV, and unclassified, suggesting higher variations among the samples within these classes. Of note, BCs associated with the four CMS classes, especially class III and IV, contain genes that highly overlap with the putative CMS marker genes; while the CMS marker genes rarely show up in BCs associated with the unclassified samples, as shown in Figure 2G. This indicates that the genes we identified in the BCs are indeed coherent with the marker genes of CMS class. Very few BCs are observed to have associations with the samples of multiple CMS classes.

Among all the BCs associated with DFS, 42.9% also over-represent certain CMS classes, while this rate is 49.5% for OS (See Figure 2H), on average. Particularly, 53.1% and 40.4% of these CMS-specific BCs fall under CMS IV class for DFS and OS respectively, on average. For DFS, the CMS IV specific BCs enrich the following pathways: glycosaminoglycan biosynthesis and metabolism, UDP glycosyltransferase, lipid, phospholipid and glycosphingolipid metabolism, mRNA splicing, and steroid hormone metabolism; while for OS, the pathways are: immune signaling, WNT and MYC signaling, VEGF signaling, tumor necrosis, notch signaling, cell proliferation and integrin pathways. This observation suggests that the extracellular matrix, glycosaminoglycan metabolism, lipid metabolism are prognostic markers for DFS if the patients are diagnosed with CMS class IV, while for OS, the markers are related to stromal infiltration. Similarly, we also observed a large proportion of CMS class I (19.1%) and CMS II specific (17.7%) BCs for DFS associated BCs, and CMS II specific (25.1%) BCs for OS associated BCs. The CMS I specific DFS associated BCs enrich chemokine signaling, integrin signaling, chondroitin sulfate and sulfur metabolism, O linked glycosylation, and other immune and inflammation related pathways; CMS II specific DFS associated BCs enrich hypoxia response, O linked glycosylation, PI3K signaling, apoptosis, and immune response pathways; and CMS II specific OS associated BCs enrich cell cycle, nucleotide excision repair, and MYC signaling pathways.

It is noteworthy that the T cell and leukocyte activation is a significant OS dependent feature for CMS1 patients but not for other CMS classes (Figure 3). CMS I has high MSI, mutation load and immune response, associated with higher abundance of neo-antigen and better response to immune-therapy (4). A high (CD8+) T-cell infiltration and activation in this group contributes to higher anti-tumor immune population. For the rest of the classes, CMS III generally has low infiltration level of T cells, and we suspect the even though cancers of CMS II and IV have high T cell infiltration, but these T cell are either exhausted or non-cancer associated. Hence the tissue level T cell gene expression do not show associations with the patients’ prognosis in any of CMS class II-IV. Such observations suggest the divergence of prognosis associated mechanisms among different CMS groups.

In addition to these, we constructed multi-variant Cox regression model to explain the patients’ prognosis using selected prognosis associated BCs and CMS class. (see Methods). Our analysis suggested that the BCs forming independent predictive markers for DFS enrich pathways including chemokine receptor, O-linked glycan biosynthesis, apoptosis, mitochondria, cell membrane, MAPK activity, tissue morphogenesis, VEGFR pathway, lipid homeostasis and cell surface receptor activity; while for OS, the BCs enrich cell death, cell proliferation, mitosis, glycosaminoglycan synthesis, integrin (possibly suggests stromal infiltration level), T cell activation, WNT beta-catenin signaling, leukocyte activation, extracellular region and glucose transport and VEGFR pathway.

In summary, our analysis reveals distinct prognosis markers of different prognosis type and CMS class. Specifically, the DFS markers are largely enriched by genes related to micro-environmental stresses while the OS markers is more determined by the level of stromal infiltration and immune response.

### Alternative drug resistance mechanisms of CRC

Chemo-therapy is one of the standard cancer treatment methods that induces cell death of fast proliferating cancer cells (20). It has been reported that cancer cells could develop resistance mechanism to chemo-therapy through alterations in pathways including cell proliferation, apoptosis, DNA damage repairing and stress response through changes in expression levels and/or mutation status of key genes (21, 22). Our understanding of drug resistance mechanism is largely complicated by intra-tumor heterogeneity within a tumor tissue and its intricate micro-environmental stresses. It is noteworthy that multiple alternative resistance mechanisms may exist among the patients, where each patient’s cancer cells acquiring one or several such mechanisms can suffer from poor prognosis to chemo-therapy. In this study, we attempt to identify the multiple chemo-resistance mechanisms within a heterogeneous patients population by our bi-clustering formulation. We hypothesize that the alternative resistance mechanisms among patients could be reflected by the BCs associated with poor prognosis to a certain chemo-drug.

The clinical information in TCGA provides patients’ treatment response to three most prevalent CRC chemo-therapy plans, including 5-Fluorouracil (5-FU), Oxaliplatin (OXA), and the combination of OXA, 5-FU and Leucovorin (FOLFOX). We selected those BCs associated with resistances to the three drugs with TCGA expression data. A BC is defined as associated with resistance of a chemo-drug if the following two conditions are both met: (1) among drug treated samples, the overall survival of samples in the BC is significantly worse than those not in the BC (p<0.001); and (2) among samples in the BC, the overall survival of samples that are drug treated is significantly worse than those not treated (p<0.05). Among the resistance associated BCs, we posit that multiple may correspond to the same resistance mechanism. In order to identify the most unique set, we incorporated a log-rank test coupled with agglomerative clustering to cluster the BCs of similar resistance mechanisms into groups, each of which is linked with one unique drug resistance mechanism (see details in Methods section).

To identify resistance mechanism associated BCs, we conducted an agglomerative clustering and log-rank test based approach to group the BCs that are highly represented by poor responders. Specifically, we generate agglomerative clustering for all the drug resistance related BCs where the distance of a pair of BCs is measured by the Jaccard index of the samples in the two BCs. Two BCs are clustered if at least one of the two sample set differences between the two BCs are insignificantly associated with drug resistance. Completed information of BC groups are given in Supplementary Table 4.

5-FU is one of the most commonly used chemo-drugs in treating CRC (23). We identified 11 BCs associated with 5FU resistance. Agglomerative clustering and stepwise test revealed that the 11 BCs form four groups, where each group consists of a number of genes tightly co-expressed, and a number of samples presented with 5FU resistance, as shown in Figure 4A. The first BC group is highly enriched by the genes involved in known chemo-resistance related mechanisms, including over expression of CFLAR involved in apoptosis and FAS signaling; CAPRIN2 related to cell proliferation and cancer multi-drug resistance; DNA excision repair gene XPA; cell cycle regulating proteins DMTF1 and SYCE2; killer cell activating receptor associated protein TYROBP; taurine metabolism gene CSAD; RNA processing proteins RBM6 and CLK1; DNA binding and transcriptional regulatory genes ZNF638, ZNF169, ZNF26, ZNF333, ZNF493, ZNF234 and ZNF33A; OGT, TAS2R5, LTB4R2 related to cellular response to chemical stimuli. It is noteworthy that a number of genes in this panel including CFLAR, CAPRIN2, XPA, TYROBP, CLK1, OGT, and LTB4R2 have been previously identified to relate to chemo-resistance in other cancer types (24–29). The second BC group is composed by highly expressed genes including SMAD2, SMAD4, TCF12, ELP2, ATG2B, PIGN, MBP, NCBP3 and PIK3C3, which enrich pathways of cell cycle, cell metabolism regulation, TGF-beta signaling, PI3K cascade, autophagy, immune responses and mRNA production regulation. The third BC group is enriched by a large number of pseudo genes and the protein coding genes in this group enrich the translation regulation and viral infection, in which genes TMA7, DEXI and EIF3CL have been previously reported as related to cisplatin and fluorouracil resistance in bladder and gastric cancer (30, 31). In addition, the four BCs group are also enriched by two different groups of ribosome proteins, which are related to translational control and elongation of peptides.

**Figure 4.**
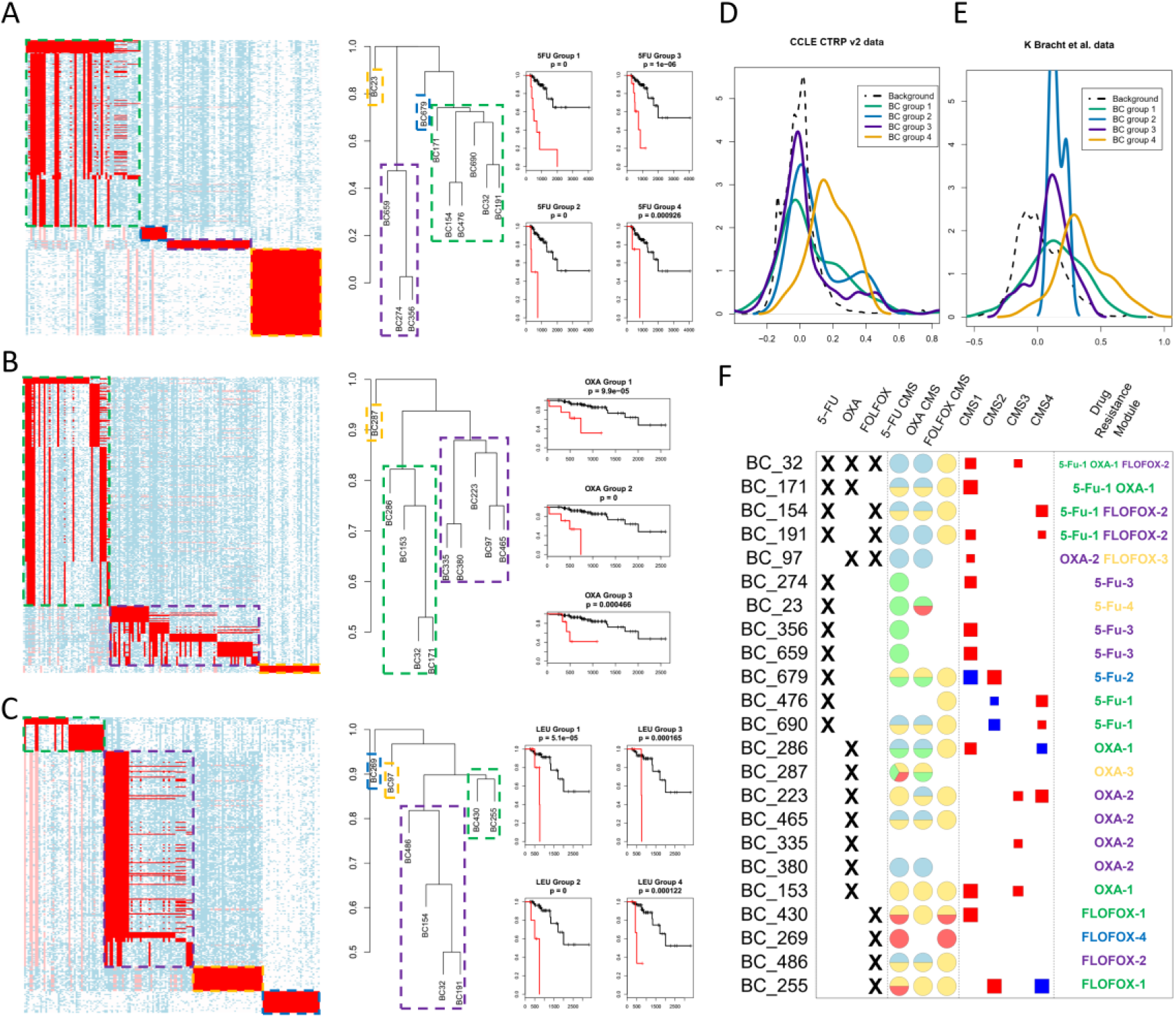
Possible alternative chemo-resistance mechanism depicted by BC groups. (A-C) Discretized gene expression profile of the resistance BC groups for 5FU (A), OXA (B), and FOLFOX (C). For (A-C), in the left-most panels, blue and white in the heatmap represent 1s and 0s in the discretized data matrix, while red represents the matrix element belonging to a certain BC group, framed in green dashed line. In the middle panels, the dendrograms show the results of agglomerative clustering of the resistance associated BCs. Each BC group is framed by a dashed rectangle. In the right-most panels, the survival curves represent for the drug treated patients, the comparison of overall survival of the patients in a BC group (red) with those not (black). (D-E) Distribution of the correlations calculated between expressions of genes in different groups with drug resistance measure IC50, in CTRP v2 dataset (D) and GI50 in K Bracht et al.’s dataset (E). The x-axis represents the correlations and the y-axis represents the density. (F) Relationships between chemo-resistance BCs and different CMS classes. In columns 1-3, a “cross” sign indicates the drugs that samples in the BCs show resistance for; in columns 4-6, larger sizes of the sectors indicate higher significances that the BC’s resistance mechanisms is also exhibited in CMS I (blue), II (yellow), III (green), and IV (red); in columns 7-10, larger sizes of the squares indicate higher significances that the BC is positively (blue)/negatively (red) enriched by samples in each CMS class (only p<0.001 are shown); the last column shows for each BC, the type of drug and BC group it is linked to.

OXA is a platinum-based antineoplastic chemo-drug used to treat colorectal cancer (23). We have identified 10 BCs with strong associations to OXA resistance, which were further clustered into three groups as shown in Figure 4B. The first BC group shows an overlap with the first group in 5FU resistance, in that the genes are also involved in known chemo-resistance related mechanisms including CFLAR, CAPRIN2, TYROBP, CLK1, OGT and LTB4R2 as well as SYCE2, RBM6, ZNF638, ZNF169, ZNF26, ZNF333, ZNF493, ZNF234 and ZNF33A related to cell cycle, mRNA processing and DNA binding. Meanwhile, this group also contains overly expressed DNA synthesis and cell cycle genes POLA1, CHFR, and TAF1; mRNA processing gene PCF11; EPHA7 and COL4A3 related to tissue development; and ITPR2 related to calcium dependent signaling transduction. The second group also contains CFLAR, CAPRIN2, SYCE2, and LTB4R2 identified in the first group. In addition, this group also contains cyclin-D binding transcription factor DMTF1; transcriptional regulation co-factor EP300; GTF2H4 related to RNA polymerase II transcription initiation; mRNA splicing gene DDX39B; and cell surface channel, transporter or exchanger genes PKD2, TRAPPC10, SMG1, and TRIO. The third group contains a number of nuclear ribonucleoproteins and HSPA5, where the latter has been previously identified as a chemo-resistance biomarker and molecular target in B-lineage acute lymphoblastic leukemia (32).

FOLFOX is combinatorial therapy of 5Fu, OXA with Leu--a reduced folic acid based drug that is used in combination with other chemotherapies to enhance effectiveness or prevent side effects of the chemo-drugs (23, 33). We have identified eight BCs forming four BC groups (Figure 4C). The first BC group shows strong overlaps with the first group of 5FU chemo-resistance, and the first and second group of OXA chemo-resistance, which includes CFLAR, CAPRIN2, SYCE2, CSAD, MSH5, XPA, OGT, LTB4R2, ZNF234, ZNF169, ZNF493, ZNF26, and ZNF333. The second group is composed of highly expressed JAK2, which is involved in multiple cytokine receptor signaling pathways related to immune response; Rho GTPase Activating Protein DLC1 (tumor suppressor); cell death related genes NME1, BCL2L15 and RPSS3A; tissue development regulating gene FOXA2; TCA cycle and respiration electron transport genes ATP5C1 and COX7A2L; and mitochondrial inner membrane translocase TIMM23. In addition, this group is also highly enriched by overly expressed ribosome proteins. The third group contains highly expressed CAPRIN2, cell proliferation regulating gene DMTF1 and mRNA processing proteins DDX39B and GTF2H4. The fourth group is composed of under expressed microRNA MIR3911 and antisense mRNA EIF1AX-AS1.

To validate the drug resistance mechanism we identified using BCs, we collected independent datasets of drug screening on colon cancer cell line (see methods). Unfortunately, to the best of our knowledge, 5-FU is the only one drug with a wide spectrum of sensitivity measure on cell lines among the three. 5-FU screening was performed on 29 and 19 colon cancer cell lines for two independent datasets (34, 35). In each dataset, we computed the correlations between the basal level expressions of all the genes and cell’s response to 5-FU, measured by IC50 and GI50 (see Supplementary table 5). Distribution of the correlations for genes in each BC group was compared with the distribution of the correlation for all genes, which serves as a random background. Density curves of the correlations of each BC group and the background are shown in Figure 4D and 4E. We have seen that, comparing with the background correlation level, genes in BC group 4 show much higher correlations to cells’ resistance to 5-Fu, and BC groups 1-3 also contain a marked portion of genes that are more correlated with 5-Fu resistance than background. This serves as further validation of our observations of alternative drug resistance mechanisms. Detailed lists of the validation data are provided in Supplementary Table 5.

In summary, for each chemo-drug, we have identified a few resistance mechanisms, some of which are novel to CRC, and they are presented in the form of BC groups. It is noteworthy that the genes CFLAR, CAPRIN2, SYCE2, OGT, and LTB4R2 are consistently observed as resistance associated for all the three drugs. Further investigation of the sample distribution of the BC groups suggests that the first BC group of 5-Fu, OXA and the second BC group of FOLFOX highly overlap, which correspond to poor response of 5-Fu and OXA in CMS1 samples and FOLFOX in CMS2 samples (Figure 4F). The second BC cluster of OXA and the third BC cluster of FOLFOX overlap, which corresponds to poor response in CMS1 samples. In addition, the 5-Fu BC groups 2, 3 and 4 show that patients of CMS III, CMS III/IV and CMS II/III are particularly resistant to 5-Fu; OXA BC groups 2 and 3 show that OXA resistance is in particular obvious in CMS II/III and CMS I/II/III; FOLFOX BC groups 1, 3, and 4 show that resistance of the drug prevalently happen to patients of CMS II/IV, CMS II and CMS IV. Interestingly, 5-Fu BC group 1 and FOLFOX BC groups 1 and 4 do not seem to show chemo-resistance mechanisms specific to any CMS classes. Among the identified BC groups for each drug type, some of them are enriched by genes involved in chemo-resistance related biological processes or known chemo-resistance markers. Meanwhile, we have seen in 1-2 BC groups for each drug type there exists novel biomarkers, including overly expressed ribosome genes and under expressed ncRNAs.

### Bi-clusters associated with mutations

We have also tested the association between BCs and 117 high frequently-mutated and non-MSI-associated genes in TCGA COAD data. Our analysis identified that 29.1% (550/1886) of the BCs annotated by the aforementioned four types of associations and 22.5% (168/746) of the unannotated BCs are associated with at least one of the gene mutations. Interestingly, among the BCs that are associated with at least one gene mutation, a large proportion of the mutations happen in genes including TMEM132D, BCL9L, NF-1, SCN10A, PCDHA10, DIP2C, GLI3, TET2, and ARFGEF2, while only a small number fall into key CRC associated gene including APC, TP53, KRAS, CTNNB1, and PIK3CA. The mutation associated BCs majorly enrich pathways of nucleotide and glucose metabolism and immune responses. Detailed pathway enrichment of each gene mutation associated BCs is given through GitHub and described in Supplementary Table 2.

## DISCUSSIONS

Disease subtype and drug therapy specific prognostic markers can offer valuable guidance in precision medicine. High throughput transcriptomics data of large cohort studies enables comprehensive identifications of prognostic markers on whole genome level. However, with patient specific features such as disease subtypes, drug treatment or other clinicopathological features, a limited number of samples is often stratified into even finer classes wherein each has a small number of samples. In such case, the statistical power on each stratified class of samples is largely reduced. Moreover, even though CMS and other cancer subtyping methods have used highly cancer relevant features, when looking at a particular drug response or prognosis, multiple alternative alterations may exist in specific but unknown subset of samples, which may or may not overlap with a certain stratification. In addition, multiple genes may interactively contribute to one response mechanism, which is especially the case in terms of drug resistance markers, as alterations in multiple pathways are always employed in one off-target resistance mechanism (36–38). How alternative drug resistance mechanisms (and their combinations) are correlated with disease subtypes or other clinicopathological features is largely undiscovered. Limiting our analysis into a pre-defining cancer subtyping or signature pathways would be a potential hurdle that could not only be misleading, but also severely harm the statistical power.

Our unsupervised bi-clustering based approach have the following advantages in identifying alternative disease subtypes/ drug therapy specific prognostic gene markers: (1) efficiently control false discoveries; (2) readily detect informative co-expressed prognostic markers; (3) conveniently handle the intricate relationships among different subtypes, and their interactions with various clinical outcomes. Of note, deriving prognostic or predictive markers from BCs with high statistical significance could not only decrease the number of independent tests but also limiting markers to co-expression gene modules, the expression level of which are more relevant in the disease context. The sample compositions in each BC provides an easily comprehensible way to understand the underlying subtypes, as well as the functional modules being executed in the BC. Our analysis has clearly demonstrated that bi-clustering based approach can effectively identify biomarkers for alternative prognosis related or drug resistance mechanisms from large scale transcriptomics data. We posit that bi-clustering is more sensitive to locate the biomarkers specific to small subset of samples and the inference on the multiple genes in the BC can be provide more biologically coherent interpretations.

Nonetheless, we have seen a few more challenges that remain to be solved beyond this study: (1) most of current bi-clustering methods tend to exclude the highly overlapping BCs, which may be problematic when consistency of BCs across different datasets are to be performed. This raises a demand for effective identification of bi-clusters with high consistency through different data sets; (2) our current analysis pipeline lacks a predicative model using BCs, which largely limits its potential of practice. A possible solution is to incorporate the bi-clusters with a binary matrix factorization formulation, i.e. treating each BC as a column basis of the discretized data matrix, and the predictive model could be built between an outcome variable and the sets of explanatory variables consisting of the loadings of all the BC bases; (3) it is noteworthy some genes within a prognosis or drug resistance predictive BC are only selected because they are co-expressed (or co-regulated) with the true prognosis or drug resistance associated genes, and the third challenge remains to identify the genes that truly contribute to the poor prognosis or drug resistance that can become possible drug targets; and (4) the BC’s statistical significance is estimated by an estimation formula for the upper bound of *p* value. The current method works well for the BCs with small number of 0s, but an improvement is need for the BCs with low consistency. We fully anticipate these challenges can be solved in future studies to increase the feasibility of BC based biomarker study.

Overall, our analysis generated a comprehensive annotation of BC based co-expression modules in CRC that offers novel biological characterizations for CMS classification and brings new insight of disease subtype and drug therapy specific prognosis predictive markers. The analysis procedures including bi-clustering formulation, identification, significance assessment and parameter settings are provided through https://github.com/changwn/BC-CRC, that can be more generally applied in precision medicine study of other disease types.

## METHODS

### Data collection

We have collected transcriptomics data of 1,440 colorectal cancer tissue samples including the one RNA-Seq data from TCGA and seven microarray data sets from GEO database. The micro-array datasets are selected with the following criteria: (1) data are collected by the top 10 most frequently utilized human microarray platforms in GEO database; (2) dataset has more than 50 samples; and (3) dataset also provide certain prognostic or clinical outcome information. We use RPKM normalized expression value for RNA-Seq data and RMA normalized expression for microarray data. Detailed data information is provided in Table 2. The DFS used in this study is defined as starting at primary treatment and stopping at disease relapse or death. Expression of each gene with multiple probes is assessed by expression of the probe with highest mean expression value in each data set. Genes of mean expressions at bottom 30% quantile in each microarray data set, and genes with 0 expression in more than 85% samples in the RNA-Seq data set are removed from the analysis, in order to control the noise of non- or lowly-expressed genes.

**Table 2.** Data information of the analyzed data.

| Data ID | Sample# | Follow-up | Platform | Normalization |
| --- | --- | --- | --- | --- |
| **GSE14333** | 290 | DFS | Affymetrix U133 Plus 2.0 | RMA |
| **GSE17536** | 177 | OS/DFS | Affymetrix U133 Plus 2.0 | RMA |
| **GSE29621** | 65 | OS/DFS | Affymetrix U133 Plus 2.0 | RMA |
| **GSE33113** | 90 | DFS | Affymetrix U133 Plus 2.0 | RMA |
| **GSE37892** | 130 | DFS | Affymetrix U133 Plus 2.0 | RMA |
| **GSE38832** | 122 | OS/DFS | Affymetrix U133 Plus 2.0 | RMA |
| **GSE39582** | 566 | OS/DFS | Affymetrix U133 Plus 2.0 | RMA |
| **TCGA-COAD** | 385 | OS | RNA-Seq | RPKM |

### Colon cancer consensus molecular subtype prediction

We applied the R package CMSclassifier to predict the CMS classification of each sample in the eight data sets (39), by which each sample will be predicted with four CMS scores representing its similarity to the four CMS classes. One sample is classified to one subtype if its CMS score of the subtype is larger than 0.5 and a sample is considered as with multiple-classification if both top two CMS scores are larger than 0.5 and the difference between the two scores is smaller than 0.1.

### Modeling the regulatory states of gene expressions via data discretization

To capture the regulatory states of a gene, we re-format the original expression data matrix into a larger binary matrix. Specifically, for a gene expression data *X*_m×n_ with m genes and n samples, we first generate a K × n binary matrix Y_g_ for each gene g. Y_g_[*i,j*] = 1 if and only if *X*[*g,j*] is in the *i*th quantile of *X*[*g*,], *i* = 1, …, *K*. Hence each row of Y_g_ indicates the samples with certain expression patterns of *g*. Then we merge all the Y_g_ to form a Km × n binary matrix *Y*_Km×n_ and apply our in-house bi-clustering software QUBIC-R to identify the bi-clusters enriched by 1s in *Y*_Km×n_.

The rationality of this formulation is that each of the bi-cluster identified here corresponds to a group of genes, the expression levels of each of which, are highly consistent over a subset of samples, hence representing a gene co-expression module specific to the subset of samples. It is worth noting that samples in one bi-cluster are highly likely to share similar transcriptional regulatory signals controlling the relevant genes. More discussion about the connection between bi-clusters and gene expression control are available in Supplementary Method.

To select a proper K, we have generated binary matrices for each data set by using K=2, 3, 4, and 5 and examined the rate of the bi-clusters that are significantly associated with (1) biological pathways, (2) clinical features, and (3) CMS classification, among all the significant bi-clusters identified in each binary matrix. On average, highest rates of significant BCs are achieved when K=3 throughout all the eight data sets (See more details in Supplementary Figure S1).

### Bi-clustering analysis of binary matrices

We applied our recently released bi-clustering R package – QUBIC-R to identify bi-clusters in discretized matrices. It is noteworthy that the number of rows ranges from 28,754 to 71,940 in this analysis. To the best of our knowledge, QUBIC R package is the most efficient bi-clustering software in the public domain that can handle input data of such large scale. The three parameters are set as follow: consistency level c=0.25, desired output number o=3000, and bicluster overlapping rate f is set at five different levels, 0.85, 0.875, 0.9, 0.95, and 1, depending on the input data size and number of 1s in each row. Detailed information for bi-clustering parameters determination and program running for each dataset are available in Supplementary Method.

By extending Xing Sun *et al*.’s work (40–42), we derived an analytical formula to estimate the upper bound of significance values for the BCs. Suppose in a random binary matrix M with *m*_0_ rows and *n*_0_ columns, its probability of 1 for any element, namely, *p*(*M*[*i,j*] = 1), is denoted as *p*_0_. Then the upper bound of the probability that at least one submatrix *M*_1_ exists in *M* could be assessed by the following formula, where *M*_1_ has *m*_1_ rows, *n*_1_ columns *z*_0_ total number of 0, and *n*_1_ ≥ *K*:

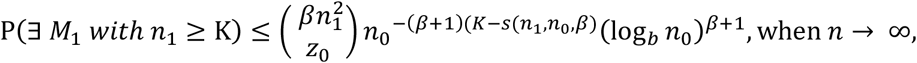

where

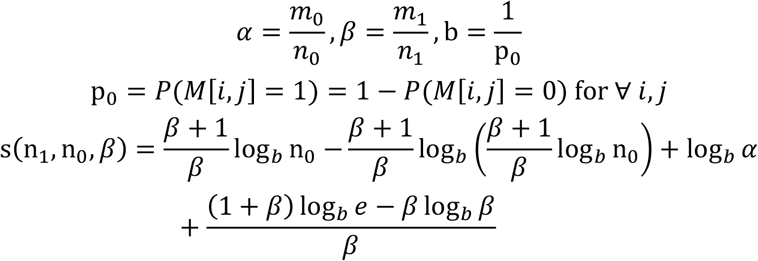

More details of the derivation of this estimation formula is given in Supplementary Method. We have tested this significance estimation method on simulated data and compared its performance with the Chernoff’s bound method (43), which is a popular measure for the effectiveness of biclustering methods. In detail, we conducted bi-clustering analysis by using same parameters on randomly generated gene expression matrices with same sizes. Significance values for the identified BCs are evaluated using both two methods and are compared with empirical *p* values. The analysis revealed that *p* values generated by our methods can more accurately recover the empirical *p* values comparing to the Chernoff’s bound method. Particularly, our method offers a good control of false discover rate for the BCs that are highly enriched by 1s, hence it is more robust in picking out the significant ones from a large number of BCs identified in a large matrix. This is particularly key to large–scale matrix.

### Annotations of the biological and clinical characteristics for each bi-cluster

Biological characteristics of each BC is assessed by whether genes in the BC significantly enrich a biology pathways or gene set. The enrichment analysis is computed by hypergeometric test, and in total, 1329 canonical gene sets including KEGG, BIOCARTA, REACTOME pathways and 1472 GO terms from MsigDB are used in the study. Here *p*=0.005 is used as the cutoff for significance.

Association analysis of each BC with clinical features were conducted using different tests based on the nature of the feature. For discrete clinical features including CMS classifications and pathological stages, we utilized fisher exact test; for continuous clinical features except for survival outcome, we compared the feature value for samples in and out of the BC by Mann Whitney test. p<0.005 is used as significance cutoff for all these tests. Notably, associations with CMS are conducted for only BCs containing more than five samples of the CMS class. For survival outcomes including DFS and OS, we compared the survival for samples in and out of the BC, using log-rank test with significance cutoff *p*<0.05.

### Analysis of somatic mutations in TCGA data

TCGA COAD level 2 mutation profile of 429 samples predicted by *mutect* is retrieved from GDC database. A total of 932 genes with mutations in more than 5% (22/429) samples are selected. Considering high MSI causes the CRC genomes to be hyper-mutated, we exclude a majority of the 932 genes whose mutations are highly associated with MSI, and 73 gene mutations not associated with MSI are retained for further analysis. The association of a gene’s mutation and MSI is calculated as the association between gene mutation and CMS class I—the class known to have high MSI, using Chi-square test (*p*<0.1).

### Multiple variable cox-regression model with variable selections

In order to identify the BCs that could best predict prognosis, we constructed multiple variable Cox-regression model between patients’ survival and the BCs shown to be associated with survival with a variable selection procedure. Here, each BC is coded into one binary explanatory vector with 1’s for samples in the BC and 0’s for samples not in the BC. Specifically, we applied forward and backward stepwise variable selection approach to select the model with lowest AIC value by using SURVIVAL and MASS package R..

### Agglomerative clustering and stepwise log-rank test based approach for identification of alternative drug resistance associated BC groups

Among the BCs that are detected to show resistance to the chemo-drugs, we posit that each BC suggests one mechanism for the drug resistance. However, there may exist more than one BC corresponding to the same mechanism. In order to identify the most unique set of resistance mechanisms, we incorporated a log-rank test coupled with agglomerative clustering to cluster the BCs of similar resistance mechanisms into groups, each of which is linked with one drug resistance.

To do this, we first defined the distance between any two BCs as 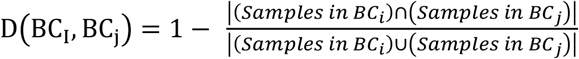, based on which an agglomerative clustering was performed. In each step of the clustering, two clusters *X* and *Y* are merged, if (1) samples in *X* ∩ *Y* is significantly associated with resistance to the drug, (2) neither samples in *X*\*Y* or *Y*\*X* is significantly associated with the drug resistance. A sample collection is defined as associated with resistance of a chemo-drug if the following two conditions are both met: (1) among drug treated samples, the overall survival of samples in the collection is significantly worse than those not in the collection (p<0.001); and (2) among samples in the collection, the overall survival of samples that are drug treated is significantly worse than those not treated (p<0.05). The agglomeration is stopped when no clusters could be merged.

## REFERENCES

1. Siegel RL, Miller KD, & Jemal A (2018) Cancer statistics, 2018. CA Cancer J Clin 68(1):7–30.

2. Wolf AMD, et al. (2018) Colorectal cancer screening for average-risk adults: 2018 guideline update from the American Cancer Society. CA Cancer J Clin 68(4):250–281.

3. Inamura K (2018) Colorectal Cancers: An Update on Their Molecular Pathology. Cancers (Basel) 10(1).

4. Guinney J, et al. (2015) The consensus molecular subtypes of colorectal cancer. Nat Med 21(11):1350–1356.

5. Cancer Genome Atlas N (2012) Comprehensive molecular characterization of human colon and rectal cancer. Nature 487(7407):330–337.

6. Roepman P, et al. (2014) Colorectal cancer intrinsic subtypes predict chemotherapy benefit, deficient mismatch repair and epithelial-to-mesenchymal transition. Int J Cancer 134(3):552–562.

7. Budinska E, et al. (2013) Gene expression patterns unveil a new level of molecular heterogeneity in colorectal cancer. J Pathol 231(1):63–76.

8. Schlicker A, et al. (2012) Subtypes of primary colorectal tumors correlate with response to targeted treatment in colorectal cell lines. BMC Med Genomics 5:66.

9. Sadanandam A, et al. (2013) A colorectal cancer classification system that associates cellular phenotype and responses to therapy. Nat Med 19(5):619–625.

10. De Sousa EMF, et al. (2013) Poor-prognosis colon cancer is defined by a molecularly distinct subtype and develops from serrated precursor lesions. Nat Med 19(5):614–618.

11. Marisa L, et al. (2013) Gene expression classification of colon cancer into molecular subtypes: characterization, validation, and prognostic value. PLoS Med 10(5):e1001453.

12. Perez-Villamil B, et al. (2012) Colon cancer molecular subtypes identified by expression profiling and associated to stroma, mucinous type and different clinical behavior. BMC Cancer 12:260.

13. Pontes B, Giraldez R, & Aguilar-Ruiz JS (2015) Biclustering on expression data: A review. J Biomed Inform 57:163–180.

14. Eren K, Deveci M, Kucuktunc O, & Catalyurek UV (2013) A comparative analysis of biclustering algorithms for gene expression data. Brief Bioinform 14(3):279–292.

15. Zhang Y, et al. (2017) QUBIC: a bioconductor package for qualitative biclustering analysis of gene co-expression data. Bioinformatics 33(3):450–452.

16. Li G, Ma Q, Tang H, Paterson AH, & Xu Y (2009) QUBIC: a qualitative biclustering algorithm for analyses of gene expression data. Nucleic Acids Res 37(15):e101.

17. Xie J, Ma A, Fennell A, Ma Q, & Zhao J (2018) It is time to apply biclustering: a comprehensive review of biclustering applications in biological and biomedical data. Brief Bioinform.

18. Subramanian A, et al. (2005) Gene set enrichment analysis: a knowledge-based approach for interpreting genome-wide expression profiles. Proc Natl Acad Sci U S A 102(43): 15545–15550.

19. Zhang C, Cao S, & Xu Y (2014) Population dynamics inside cancer biomass driven by repeated hypoxia-reoxygenation cycles. Quantitative Biology 2(3):85–99.

20. DeVita VT, Jr. & Chu E (2008) A history of cancer chemotherapy. Cancer Res 68(21):8643–8653.

21. Abdullah LN & Chow EK (2013) Mechanisms of chemoresistance in cancer stem cells. Clin Transl Med 2(1):3.

22. Zheng HC (2017) The molecular mechanisms of chemoresistance in cancers. Oncotarget 8(35):59950–59964.

23. Gustavsson B, et al. (2015) A review of the evolution of systemic chemotherapy in the management of colorectal cancer. Clin Colorectal Cancer 14(1): 1–10.

24. Fraser M, et al. (2003) Chemoresistance in human ovarian cancer: the role of apoptotic regulators. Reprod Biol Endocrinol 1:66.

25. Weaver DA, et al. (2005) ABCC5, ERCC2, XPA and XRCC1 transcript abundance levels correlate with cisplatin chemoresistance in non-small cell lung cancer cell lines. Mol Cancer 4(1):18.

26. Mochmann LH, et al. (2014) ERG induces a mesenchymal-like state associated with chemoresistance in leukemia cells. Oncotarget 5(2):351–362.

27. Zhang L, et al. (2017) Clk1-regulated aerobic glycolysis is involved in glioma chemoresistance. J Neurochem 142(4):574–588.

28. Cheng S, et al. (2017) GNB2L1 and its O-GlcNAcylation regulates metastasis via modulating epithelial-mesenchymal transition in the chemoresistance of gastric cancer. PLoS One 12(8):e0182696.

29. Park J, Park SY, & Kim JH (2016) Leukotriene B4 receptor-2 contributes to chemoresistance of SK-OV-3 ovarian cancer cells through activation of signal transducer and activator of transcription-3-linked cascade. Biochim Biophys Acta 1863(2):236–243.

30. Tanaka N, et al. (2018) Single-cell RNA-seq analysis reveals the platinum resistance gene COX7B and the surrogate marker CD63. Cancer Med 7(12):6193–6204.

31. Kim M, et al. (2017) GFRA1 promotes cisplatin-induced chemoresistance in osteosarcoma by inducing autophagy. Autophagy 13(1): 149–168.

32. Uckun FM, et al. (2011) Inducing apoptosis in chemotherapy-resistant B-lineage acute lymphoblastic leukaemia cells by targeting HSPA5, a master regulator of the anti-apoptotic unfolded protein response signalling network. Br J Haematol 153(6):741–752.

33. Tsai YJ, et al. (2016) Adjuvant FOLFOX treatment for stage III colon cancer: how many cycles are enough? Springerplus 5(1):1318.

34. Rees MG, et al. (2016) Correlating chemical sensitivity and basal gene expression reveals mechanism of action. Nat Chem Biol 12(2):109–116.

35. Bracht K, Nicholls AM, Liu Y, & Bodmer WF (2010) 5-Fluorouracil response in a large panel of colorectal cancer cell lines is associated with mismatch repair deficiency. Br J Cancer 103(3):340–346.

36. Chang RL, Xie L, Xie L, Bourne PE, & Palsson BO (2010) Drug off-target effects predicted using structural analysis in the context of a metabolic network model. PLoS Comput Biol 6(9):e1000938.

37. Schenone M, Dancik V, Wagner BK, & Clemons PA (2013) Target identification and mechanism of action in chemical biology and drug discovery. Nat Chem Biol 9(4):232–240.

38. Mansoori B, Mohammadi A, Davudian S, Shirjang S, & Baradaran B (2017) The Different Mechanisms of Cancer Drug Resistance: A Brief Review. Adv Pharm Bull 7(3):339–348.

39. Eide PW, Bruun J, Lothe RA, & Sveen A (2017) CMScaller: an R package for consensus molecular subtyping of colorectal cancer pre-clinical models. Sci Rep 7(1): 16618.

40. Sun X (2007) Significance and recovery of blocks structures in binary and real-valued matrices with noise (The University of North Carolina at Chapel Hill).

41. Sun X & Nobel A (2006) Significance and recovery of block structures in binary matrices with noise. International Conference on Computational Learning Theory, (Springer), pp 109–122.

42. Sun X & Nobel AB (2008) On the size and recovery of submatrices of ones in a random binary matrix. Journal of Machine Learning Research 9(Nov):2431–2453.

43. Hoeffding W (1963) Probability Inequalities for Sums of Bounded Random Variables. Journal of the American Statistical Association 58(301): 13–30.

